# Self-regulation of the nuclear pore complex enables clogging-free crowded transport

**DOI:** 10.1101/2022.08.19.504598

**Authors:** Tiantian Zheng, Anton Zilman

## Abstract

Nuclear pore complexes (NPCs) are the main conduits for macromolecular transport into and out of the nucleus of eukaryotic cells. The central component of the NPC transport mechanism is an assembly of intrinsically disordered proteins (IDPs) that fills the NPC channel. The channel interior is further crowded by large numbers of simultaneously translocating cargo-carrying and free transport proteins. How the NPC can efficiently, rapidly and selectively transport varied cargoes in such crowded conditions remains ill understood. Past experimental results suggest that the NPC is surprisingly resistant to clogging and that transport may even become faster and more efficient as the concentration of transport protein increases. To understand the mechanisms behind these puzzling observations, we construct a computational model of the NPC comprising only a minimal set of commonly-accepted consensus features. This model qualitatively reproduces the previous experimental results and identifies self-regulating mechanisms that relieve crowding. We show that some of the crowding-alleviating mechanisms – such as preventing saturation of the bulk flux – are “robust” and rely on very general properties of crowded dynamics in confined channels, pertaining to a broad class of selective transport nanopores. By contrast, the counter-intuitive ability of the NPC to leverage crowding to achieve more efficient single molecule translocation is “fine-tuned” and relies on the particular spatial architecture of the IDP assembly in the NPC channel.

---

Nuclear pore complexes (NPCs) are macromolecular “machines” that are the main conduits of molecular transport between the nucleus and cytoplasm in eukaryotic cells and facilitate the rapid and selective transport of macromolecules between these cellular compartments [1]. NPCs are vital to all eukaryotic lifeforms and disruptions of their ability to carry out nucleocytoplasmic transport often result in severe consequences for the entire organism. Disruptions of NPC transport and alterations of NPC components have been associated with multiple cancers and neurodegenerative diseases [2–7].

The structure of the NPC is unique among cellular transporters. Scaffold nucleoporins (nups) stabilize an hourglass-shaped channel across the nuclear envelope, with an approximate average diameter of 35–50 nm [1, 8]. Multiple copies of around 10–15 different types of intrinsically disordered proteins, known as FG nucleoporins (FG nups) due to the many copies of phenylalanine and glycine repeats (FG repeats) in their sequences, are anchored to the interior of this channel [1,9]. These disordered FG nups form an assembly which occupies much of the NPC channel, and are a vital component of the NPC’s selectivity mechanism. To a large degree, FG nups behave as flexible cohesive polymers with some cross-linking due to mostly attractive hydrophobic interactions between FG repeats [1, 10–12]. The spatial and temporal organization of FG nups remains imperfectly understood, although a consensus exists that the FG assembly has a heterogeneous density profile. In the main, the FG nups anchored closer to the center of the channel are more cohesive and have stronger inter- and intra-chain attraction (more globule-like in the coil-globule paradigm of polymer physics) than the FG nups anchored near the nuclear and cytoplasmic exits (which are less cohesive and more coil-like) [10, 12–18].

The FG assembly plays a central role in the selectivity mechanism of the NPC as it forms the selective barrier for transport between the nucleus and cytoplasm. It excludes passively diffusing macromolecules in a size dependent manner, with translocation through the NPC becoming increasingly obstructed with larger molecule size [19–21]. This exclusion predominantly arises from a combination of entropic repulsion due to the displacement of thermally fluctuating FG nup chains and the cost of disrupting the cohesive FG-FG contacts between the FG nups [1, 13, 22–24]. Facilitated transport through the NPC is mediated by transport proteins known as nuclear transport receptors (NTRs). During nuclear import, cargoes in the cytoplasm carrying a nuclear localization sequence are recognized and bound by NTRs, forming an NTR-cargo complex. Attractive binding interactions between FG nups and NTRs allow the NTR-cargo complex to penetrate and translocate through the FG nup assembly [1, 13, 22, 23]. When the NTR-cargo complex reaches the nucleus, the nuclear factor RanGTP binds the NTR, thereby releasing the cargo from the complex and sequestering it within the nucleus. As the translocation of the NTR-cargo complex through the NPC is not directly coupled to GTP hydrolysis, the translocation of NTRs/NTR-cargo complexes through the NPC is a stochastic diffusive process as reflected in the common occurrence of abortive translocations, where the import complex returns to the cytoplasm intact [8, 25, 26]. Translocation of an individual NTR-cargo complex does not induce an observable conformational change between an “open” and a “closed” state of the NPC [8]; rather, multiple NTR-cargo complexes and free NTRs are present simultaneously within the FG nup assembly, and many translocation events occur in parallel [25, 27, 28]. According to previous estimates, at least 40% of the space in the NPC channel is occupied by FG nups and NTRs [29]. Therefore, a key question is how nucleocytoplasmic transport can occur rapidly and efficiently through such a crowded medium without slowing down and clogging.

Several strands of experimental evidence converge towards a surprising conclusion that the NPC does not clog with an increase in the accumulation of NTRs and small NTR-cargo complexes within the pore. On the contrary, this molecular crowding appears to lead to faster and more efficient transport. As the cargoes used in these studies are much smaller than NTRs, we will henceforth refer to NTR-cargo complexes also as NTRs, keeping in mind that these NTRs may sometimes be transporting small cargoes.

A series of experiments both in permeabilized [30] and intact [31] cells found that the flux of NTRs into the nucleus increased linearly with the cytoplasmic NTR concentration, with no indication of saturation to a plateau (Figure 2A). Flux saturation was absent even at very high concentrations of tens of *µ*M that are orders of magnitude higher than the typical measured equilibrium dissociation constants of NTR interactions with the FG assembly [32, 33]. Even more counter-intuitively, a single-molecule study found that increasing the concentration of NTRs in the cytoplasm increased their probability of translocating successfully into the nucleus while the mean translocation time decreased (Figure 2B) [26].

Thus, the indicative features of clogging resulting from channel obstruction – flux saturation, a reduced probability of successful translocations and slowing down of the transport times – appear absent from these measurements [33–35].

An additional longstanding experimental puzzle is the observation of a long-lived fraction of endogenous NTRs within permeabilized cells, which leave the NPC at timescales (minutes and hours) that are much greater than expected from single molecule transport times (milliseconds) [36–39]. This phenomenon has been reproduced in *in vitro* assemblies of surface grafted FG nups and artificial nanopores [32, 40], leading to the proposal of a “kap-centric model”, which suggests that NTRs (known also as karyopherins or “kaps”) actively control the kinetic properties of nucleocytoplasmic transport. It has been suggested that the observations of clogging-free transport [26, 31] are examples of NTR-mediated control of transport [36]. However, a rigorous theoretical explanation of the linkage between the mechanisms behind all these disparate phenomena is still missing.

In this work, we employ a minimal coarse-grained computational model of NPC transport. The model semi-quantitatively recreates the experimental observations of clogging-free transport, as well as the appearance of the “slow” and the “fast” NTR fractions. We identify the minimal set of mechanisms and structural features responsible for these observations, and explain the common underlying principles of clogging-free transport.

## 1 Model

Growing evidence indicates that despite its structural complexity, many important NPC functions can largely be understood through a small number of basic principles. A wide range of experimental observations have been reproduced using minimal models that subsume the exact molecular sequences of individual FG nups and the complex interactions taking place within the entire FG nup assembly into a small number of coarse-grained variables [11, 19, 37, 41–44]. The salient predictions of these minimal models agree in the main with those of more detailed models that take into account some aspects of FG nup sequence and NTR patterning [45,46]. In this work, we focus on these essential components of the NPC mechanism while searching for an explanation of the regulation of transport in the presence of crowding. We construct a coarse-grained computational model of the NPC with a minimal number of features (Figure 1A shows a sample snapshot), composed of a pore containing grafted polymers, representing FG nups, and spherical particles of appropriate size representing NTRs (see e.g. [19, 45, 47, 48].

**Figure 1:**
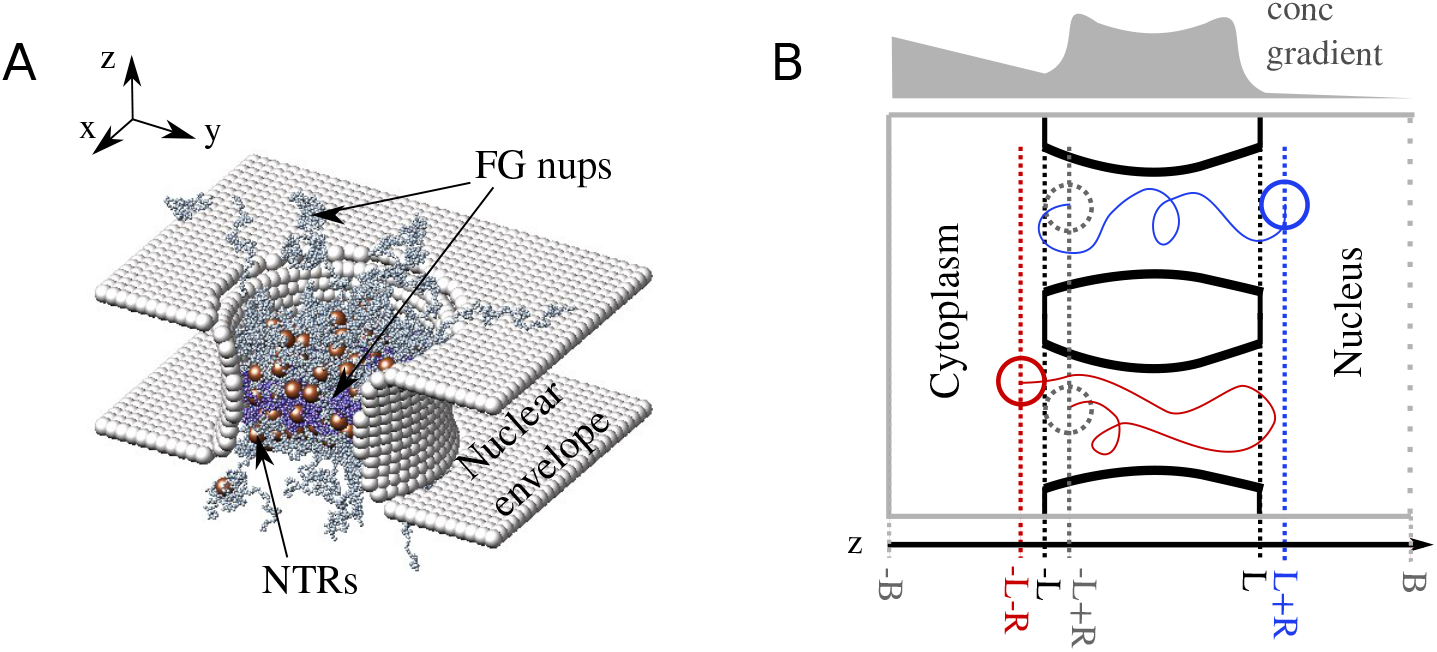
Computational model setup. (A) A snapshot from the simulation of the model of the NPC. The nuclear envelope and structural nucleoporins are represented as rigid surfaces (light grey). Two types of FG nups are grafted with 8-fold rotational symmetry to the interior of the channel, with more cohesive FG nups in the center of the NPC (purple) and less cohesive FG nups at the peripheries (grey). NTRs are modeled as spheres (brown) that interact attractively with FG nups and through steric repulsion with each other. (B) An illustration (not to scale) of the different paths followed by the center of an NTR during an entry event (blue) and an abortive event (red). We consider the NTR to have entered the pore when it crosses into the coordinate *z*_ent_ = −*L* + *R* (i.e. the entire volume of the NTR is contained within the NPC channel). The translocation ends once the entire volume of the NTR leaves the NPC channel, and resides either at 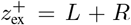 (classified as an entry event) or at 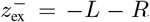 (classified as an abortive event). Simulation box boundaries are indicated in grey, the absorbing boundary (see text) is indicated by a dashed line. The length of the NPC channel is 2*L* = 40 nm and *R* = 2.5 nm. The total length of the simulation box is 120 nm. An illustrative non-equilibrium concentration profile of the NTRs is shown in light grey indicating the direction of the NTR concentration gradient; see text.

As in previous work [11, 45, 49, 50] each monomer on an FG nup has a diameter of 1 nm, and represents approximately four amino acids, corresponding to the typical size of an FG repeat. The monomers on the FG nups interact attractively with each other through a cohesive interaction, and also interact attractively with NTRs. NTRs are modelled as spheres with diameters of 5 nm and volume *v* = 65.4 nm^3^. The NPC channel is modelled as a rigid hourglass shaped structure with minimal and maximal pore diameters of 50 nm and 60 nm and length of 40 nm, as illustrated in Figure 1. FG nups are grafted along 8 spokes inside the channel, with each spoke consisting of 14 FG nups of 200 monomers each grafted uniformly along the length of the channel. Using the average molecular weight for an amino acid, this results in approximately 9 MDa of FG nups within the pore, which is typical for yeast NPCs [9]. After performing sensitivity analysis of the model using several similar pore designs, we have found the main conclusions of this paper are not sensitive to the exact choice of the model parameters.

As described previously, a finitely extensible nonlinear elastic (FENE) spring potential is applied on neighbouring monomers on the same chain to represent the bonds forming the backbone of the polymers, 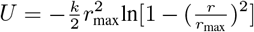, where *r*_max_ is the maximum extension of the bond, taken to be 1.5*b*, and *k* is the stiffness of the bond, taken to be 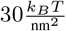 [45, 50, 51].

Interactions between all particles are modelled by a shifted Lennard-Jones type potential [52]:

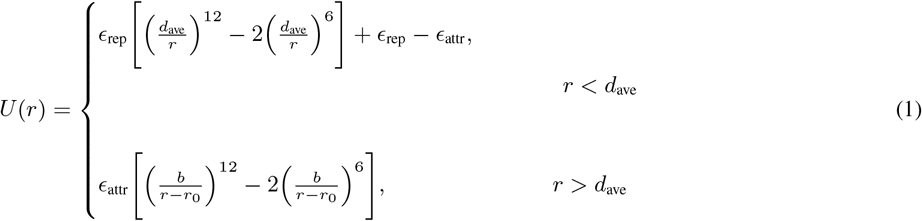

where *b* is the monomer diameter, *d*_ave_ is the average diameter of the two interacting particles, 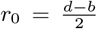 (where *d* is the cargo diameter), and *r* is the distance between the particles. The shift by *r*_0_ accounts for the difference in particle sizes, and keeps the range of interaction independent of *ϵ*_attr_. *ϵ*_rep_ and *ϵ*_attr_ are the strengths of the repulsive and attractive interactions, respectively; *ϵ*_attr_ *>* 0 for cohesive interactions between monomers and monomers (*ϵ*_coh_); and the binding interactions between monomers and NTRs (*ϵ*_NTR_).

Experimental observations indicate that the types of FG nups grafted near the centre of the NPC tend to adopt collapsed-coil configurations in solution, while the types of FG nups located near the openings of the pore tend to adopt relaxed or extended-coil conformations, corresponding to higher and lower cohesiveness respectively [10, 12–14, 16–18]. Thus, in our model, the 6 central FG nups within each spoke were modelled with a stronger cohesiveness (*ϵ*_coh_ = 0.5*k*_*B*_*T*) than the two sets of 4 FG nups located closer to the channel exits, (*ϵ*_coh_ = 0.3*k*_*B*_*T*), see Figure 1. These values of *ϵ*_coh_ were chosen to lie on either side of the coil-globule transition, representing the different chain conformations of extended and collapsed-coil FG nups [10,11] (see Supplementary Figure S1).

The interaction energy between the monomers and the NTRs was chosen phenomenologically to reproduce the levels of crowding observed experimentally within NPCs. Based on observations that there are ∼ 100 Imp*β* inside the NPC at physiological conditions [25,39], we set *ϵ*_NTR_ to be 0.95 *k*_*B*_*T* which ensures that the number of the NTRs inside the pore was between 100 – 250 at an external concentration of 1 – 10 *µM*. Using this value produces simulated equilibrium dissociation constants between NTRs and the NPC (Supplementary Figure 2) which are well aligned with experimentally measured values [53].

In order to generate a steady-state flux of NTRs through the NPC which mimics that produced by the RanGTP cycle, NTRs that reach the nuclear end of the simulation box are absorbed and “recycled” into the cytoplasmic box, while the cytoplasmic boundary is reflective (Figure 1B). This setup maintains a steady state non-equilibrium gradient of NTR concentration between the cytoplasmic and the nuclear compartments, and a resulting constant steady-state flux of NTRs in the cytoplasmic-to-nuclear direction.

The model was simulated using Brownian dynamics with an implicit solvent and a Langevin thermostat using the molecular dynamics package LAMMPS [54] using the resources provided by ComputeCanada. The time scales of our simulation were fixed by assigning the viscosity in our simulations to be the viscosity in the cytoplasm (taken as 5cP [55]), so that one timestep of our simulation corresponds to 1.35 × 10^*−*7^ ms.

## 2 Results

Our model recapitulates the experimental observations of the dynamics of NPC transport in bulk and single-molecule studies of transport [26, 30, 31].

As mentioned in Methods, within our simulation, the conditions mimicking the initial stages of nuclear import are created by continuously removing NTRs from the nuclear compartment through the absorbing boundary (see Methods and Figure 2B), which creates a non-equilibrium flux of NTRs into an empty nucleus.

**Figure 2:**
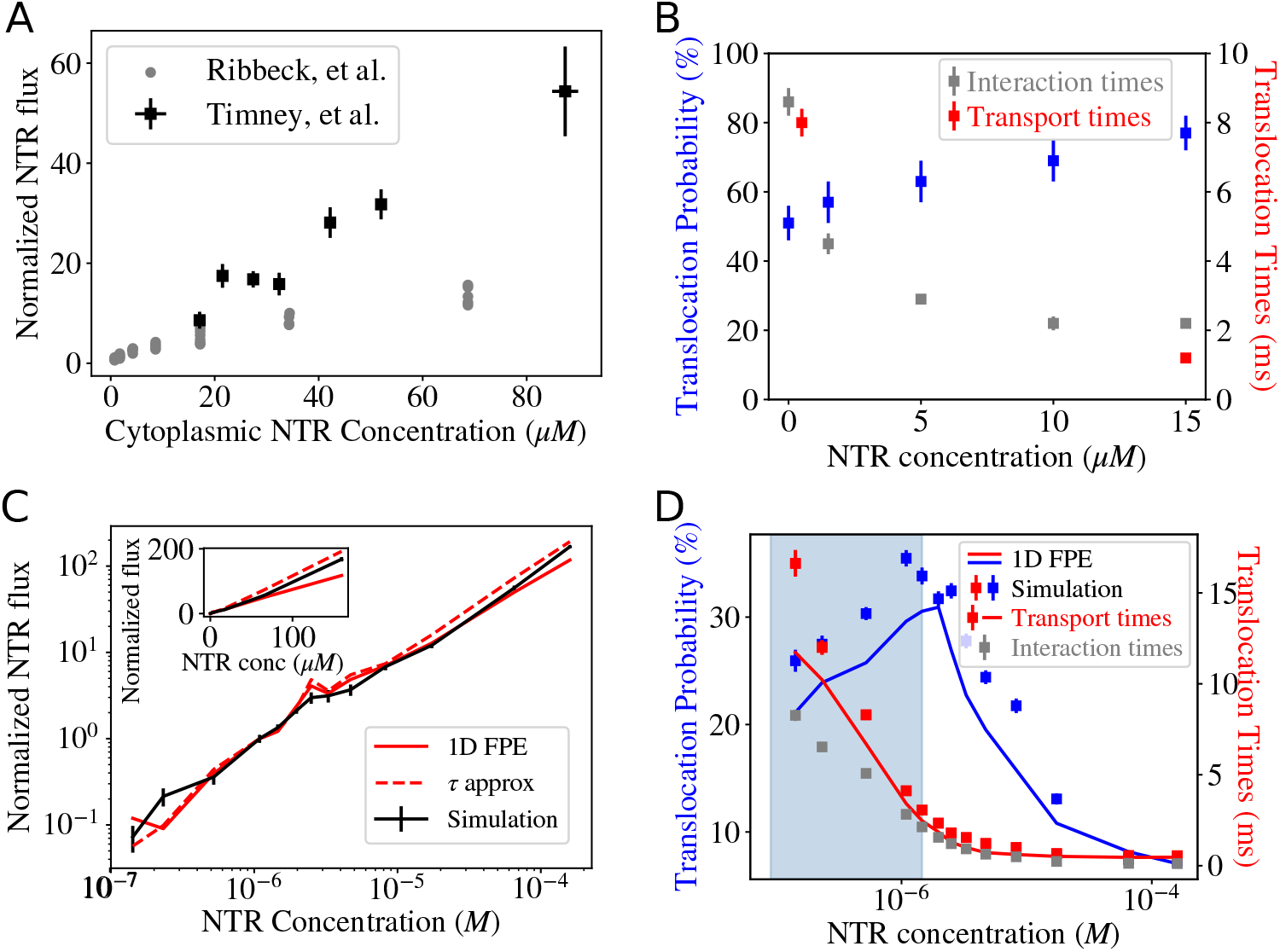
Comparison of the model predictions with the experimental data. (A) Experimentally measured flux as a function of concentration. Grey: flux of (the NTR) Transportin through the NPC (normalized by the value at 1 *µM* Transportin concentration). Adapted from [30]. Black: flux (normalized by the value at 1 *µM*) of NLS-GFP cargoes transported primarily by Kap 123p (an NTR) into the nucleus in live cells. Adapted from [31]. Both experiments show no obvious saturation of the flux into the nucleus towards a maximal value as the NTR concentration increases. (B) Single-molecule data depicting translocation probabilities which increase and transport/interaction times (see text for definitions) which decrease with increased crowding; adapted from [26]. (C) Calculated flux (normalized by the value at 1 *µM* NTR concentration) into the nucleus as a function of NTR concentration within the cytoplasm. Flux appears to increase linearly with NTR concentration, with no saturation even at very high concentrations. Black: fluxes measured from simulations, error bars indicate 1 SE; red (solid): analytical approximation using the dimensionally reduced 1D Fokker-Planck Equation diffusion model; see text and Equation 6; red (dashed): the alternative approximation using Equation 7. Inset shows the same data plotted on linear axes. (D) Simulated translocation probabilities (blue squares) and transport/interaction times (red/grey squares) as functions of the NTR concentration outside the pore. The blue shaded region indicates the regime where translocation probabilities increase and transport times decrease with crowding. Error bars indicate 1 SE. The simulation data can be explained using a dimensionally reduced 1D Fokker-Planck Equation diffusion model (solid lines); see text and Equation 6.

The main single-molecule measurements are the *translocation probability* and the *transport time*. The translocation probability is experimentally defined as the fraction of trajectories where an NTR that enters the pore on the cytoplasmic side exits the pore from the nuclear side (resulting in an “entry event”), out of all trajectories in which an NTR enters the pore (which also includes “abortive events” where the NTR leaves the pore from the cytoplasmic side) [26]. The transport time is defined as the average duration of successful entry events.

Yang et al [26] report the crowding associated decrease in the transport times – the average times of successful entry events (Figure 2B, red) – as well as a decrease in a related quantity – interaction times (Figure 2B, grey). The interaction time is defined to be the average duration of both entry and abortive events. Our model captures the decrease in both transport and interaction times; in our analysis and interpretation, we focus on the former, as a decrease in transport times is a direct indication of the absence of clogging.

In our model, NTRs are considered to have entered the pore when the entire NTR volume is contained in the NPC channel between the membranes of the nuclear envelope (NE), and are considered to have left the pore when no portion of the NTR volume is contained between the NE membranes, as shown in Figure 1B. Our definition parallels that used in [26], which considered NTRs to be inside the NPC if they reached the tips of the cytoplasmic filaments (i.e. came within 100 nm of the central plane of the NE, based on the size of human NPCs). As our NPC mimic is modelled on the smaller yeast NPC, we consider the openings of the NPC channel to be located 20 nm away from the center plane of the NE. The qualitative results of the model are not sensitive to the exact definitions of these locations (see Supplementary Information Section 1A).

As shown in Figure 2, in our model increased crowding does not result in the saturation of the flux to a plateau (Figure 2C), even for concentrations much higher than the equilibrium dissociation constant between NTRs and the NPC measured from our simulation (see Supplementary Figure S2). Clogging is absent despite the fact that up to 60% of the available volume within parts of the NPC becomes occupied by NTRs and FG nups at around 100 *µM* (Supplementary Figure S3A). The translocation times decrease with crowding (Figure 2D), and the translocation probability is non-monotonic with the NTR concentration – contrary to naive theoretical expectations [33, 56]. Most surprisingly, in a regime that spans over an order of magnitude of NTR concentrations (highlighted in blue-grey, Figure 2D), crowding increases translocation probabilities, paralleling experimental observations [26]. We refer to this regime of simultaneously increasing translocation probabilities and decreasing transport times as *optimal single-molecule transport*.

These results of the model reproduce the surprising absence of clogging in response to crowding observed in the experimental studies both on the single molecule and bulk flux levels [26, 30, 31]. In the remainder of this section, we elucidate the physical mechanisms that underpin these effects of crowding on transport, and establish the minimal set of NPC features necessary for this counter-intuitive behaviour.

### 2.1 Dimensionality reduction and effective medium approximation

In order to systematically investigate and disambiguate specific mechanisms responsible for the unusual crowding response of the NPC, we have developed a dimensionally reduced analytical approximation to the simulation models. Using an effective-medium type approach, we mapped NTR transport though the NPC onto a description as diffusion in an effective 1D potential that subsumes the many-body NTR-FG nup and NTR-NTR interactions and entropic contributions that define the free energy landscape in the pore (Equation 2). The effective potential within the pore arises through an interplay between FG nup density, FG nup cohesiveness, and the NTR-FG nup interaction energy. In our simple model, a higher density of FG nups corresponds to a deeper effective potential (Supplementary Figure S5A). The effects of crowding on transport enter through the crowding dependent changes of the 1D *effective potential* and the 1D *effective diffusion coefficient* within the channel. The effective potential and effective diffusion coefficient can both be estimated independently from auxiliary simulations carried out in equilibrium conditions (see Supplementary Information Section 1B for descriptions), and the results are shown in Figure 3. Mathematically, the transport can be described by a 1D Fokker-Planck equation [14, 57–61],

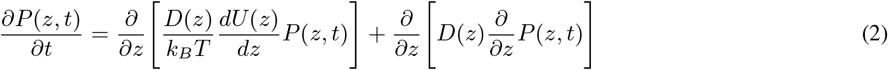

where *P*(*z, t*) = *∫*_*A*(*z*)_ *P*(*x, y, z*)*dxdy* is the number of NTRs within a slice *dz* of the pore, and *A*(*z*) is the cross sectional area of the simulation box at position *z* [59–61]. *D*(*z*) is the 1D effective diffusion coefficient at the position *z*; *U* (*z*) = − ln *P*_eq_(*z*) (see Supplementary Information Section 1B) is the effective potential at the position *z* in units of *k*_*B*_*T* [59–61]. From the Fokker-Planck equation, the translocation probability can be calculated as (see Figure 1B for definitions of *z*_ent_ and *z*_ex_):

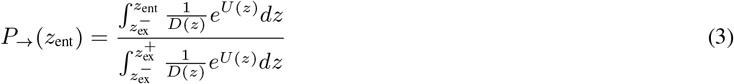

**Figure 3:**
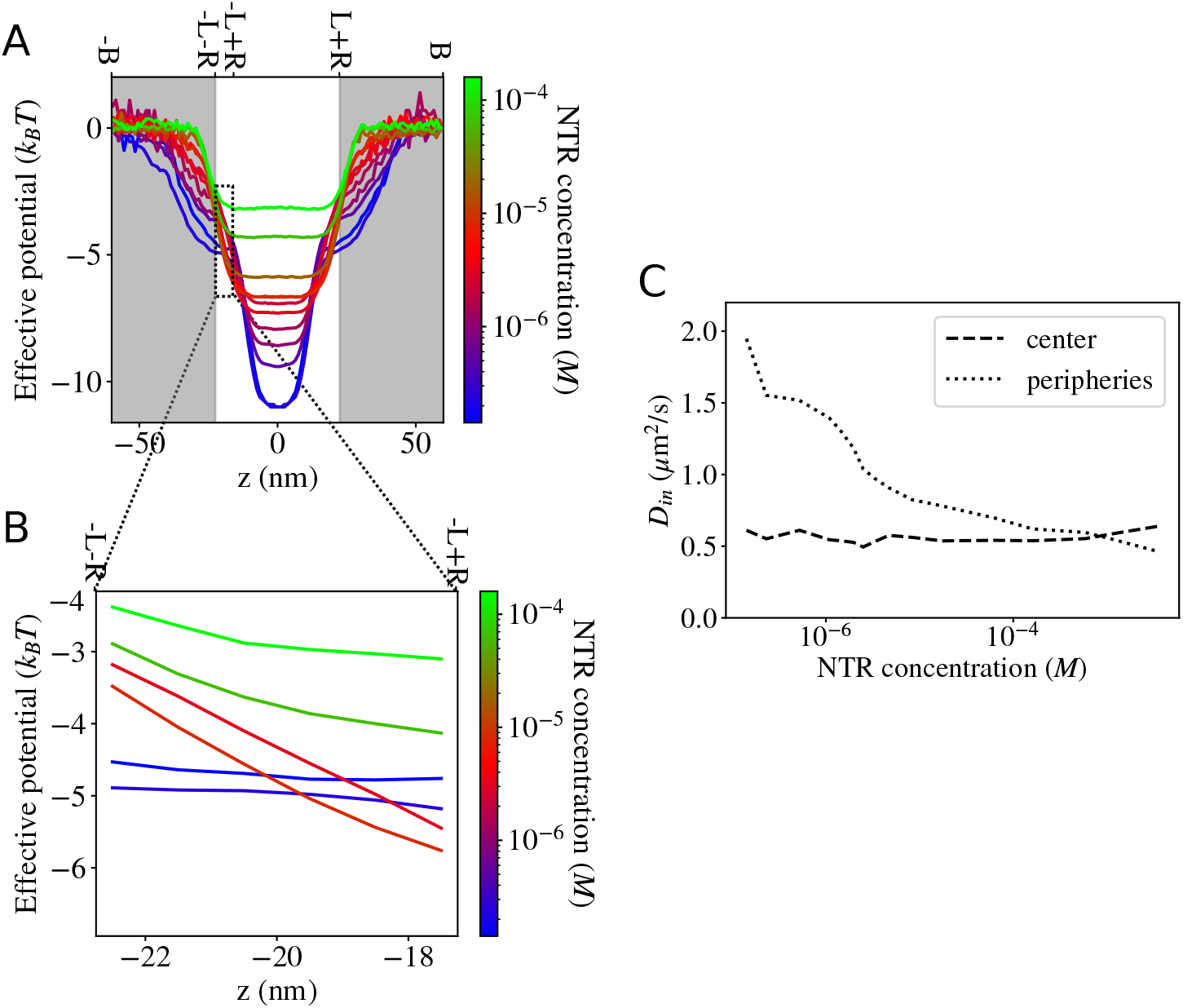
The effects of increased crowding on the effective potential and the diffusion coefficient in the pore. (A) Effective potentials measured within the simulation box. The white background indicates the region where an NTR is considered to be within the pore (for the purpose of classifying abortive and entry events). See Supplementary Information Section 1B for how effective potentials are measured in the presence of crowding. (B) Zoomed in effective potentials for representative NTR concentrations in the region from *z* = −*L* − *R* to *z* = −*L* + *R* (the bounds of the integral in the numerator of Equation 3). (C) The effective diffusion coefficients of NTRs in the central and peripheral regions as functions of the NTR concentration (see Supplementary Figure S5A).

The transport time of an entry event (the mean first passage time conditioned on exit into the nuclear compartment) [34, 57, 58] is given by

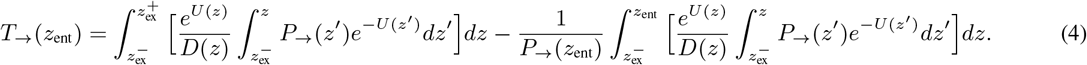

It is not obvious *a priori* that the complicated interactions and dynamics of NTR passage through the FG nup assembly and the effects of crowding can be subsumed separately into the 1D effective potential and diffusion coefficient. Nevertheless, Figure 2D shows that the results of the 1D reduced model agree well with the direct measurements from the simulations, confirming that the effective medium approximation captures the essential properties of NTR translocation even in the presence of crowding.

The 1D diffusion model also agrees well with the simulations of the bulk flux, including the non-saturating behaviour (Figure 2C, solid red line). Within the effective medium approximation, the 1D NTR density *N* (*z, t*) within the channel is given by *N* (*z, t*) = *P* (*z, t*)*N* where *N* is the total number of NTR particles in the simulation box. Thus, it is described by the same equation as Equation 2 for *P* (*z, t*). Namely, 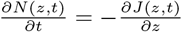, where the bulk flux of the particles is

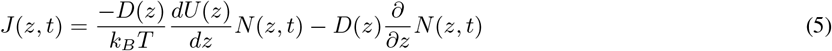

At steady-state, *J*(*z*) is constant for all *z*. Solving Equation 5, with the reflective boundary condition at −*B* (the cytoplasmic edge of the simulation box) and absorbing boundary at *B* (the farthest edge of the simulation box on the nuclear side) *N*(−*B*) = *NP*(−*B*) and *N*(*B*) = 0 (see Figure 1B), gives

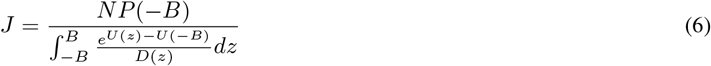

The mapping of the transport onto a 1D diffusion model that captures the simulations results (Figure 2C and D), enables us to investigate how exactly the crowding controls the transport through its effects on the effective potential and the effective diffusion coefficients. As shown in Figure 3, increased crowding produces three distinct effects. First, increased crowding decreases the average depth of the effective potential within the simulation box (Figure 3A). Second, increased crowding produces a non-monotonic and non-uniform change in the effective potential in the region between *z* = −*L* − *R* and *z* = −*L* + *R*, which corresponds to the bounds of the integral in the numerator of Equation 3 (Figure 3B). Third, increased crowding decreases the effective diffusion coefficient in the peripheries of the pore (Figure 3B).

The diffusion coefficients in our simulations decrease or remain constant with increased crowding, which, on their own, would lead to constant or longer escape times from the pore. The effect of crowding on the diffusion coefficients therefore cannot be responsible for either non-saturating flux or optimal single-molecule transport (see Supplementary Information Section 1C for the full discussion). Instead, both phenomena occur as a result of the changes to the effective potential.

In the next three subsections, we investigate how the crowding-induced changes to the effective potential profile prevent clogging and produce optimal single-molecule transport, and identify the components of the NPC architecture responsible for these effects.

### 2.2 Inter-NTR competition for space due to steric repulsion in the pore results in faster transport times and non-saturating flux

Both the computational model of the NPC and the 1D effective diffusion model reproduce the experimentally observed absence of saturation of the bulk flux shown in Figure 2C, as well as the experimentally observed decrease in the translocation times with crowding, shown in Figure 2D. These two effects are accompanied by the effective potential experienced by each NTR becoming shallower at the center of the pore with increasing NTR concentration, as shown in Figure 3A. The shallowing of the effective potential due to crowding occurs partly due to steric competition between the NTRs and is consistent with previous reports of “negative cooperativity” of NTR insertion into equilibrium surface grafted assemblies of FG nups [42, 44]. As we show in this section, this competition-driven shallowing of the effective potential is sufficient to account for the absence of clogging in NPC transport.

To establish the minimal features needed for clogging-free flux, we simulated NTRs in an FG nup-less pore, where their interactions with the FG nups were replaced by an externally applied attractive potential *W* (*z*) acting on all NTRs within the pore that corresponds to the effective potential measured in pore simulations: *W* (*z*) = *U* (*z*)+ln *A*(*x*) [59–61]. As shown in Figure 4A and B this minimal setup reproduced both the non-saturating flux and the shorter transport times.

**Figure 4:**
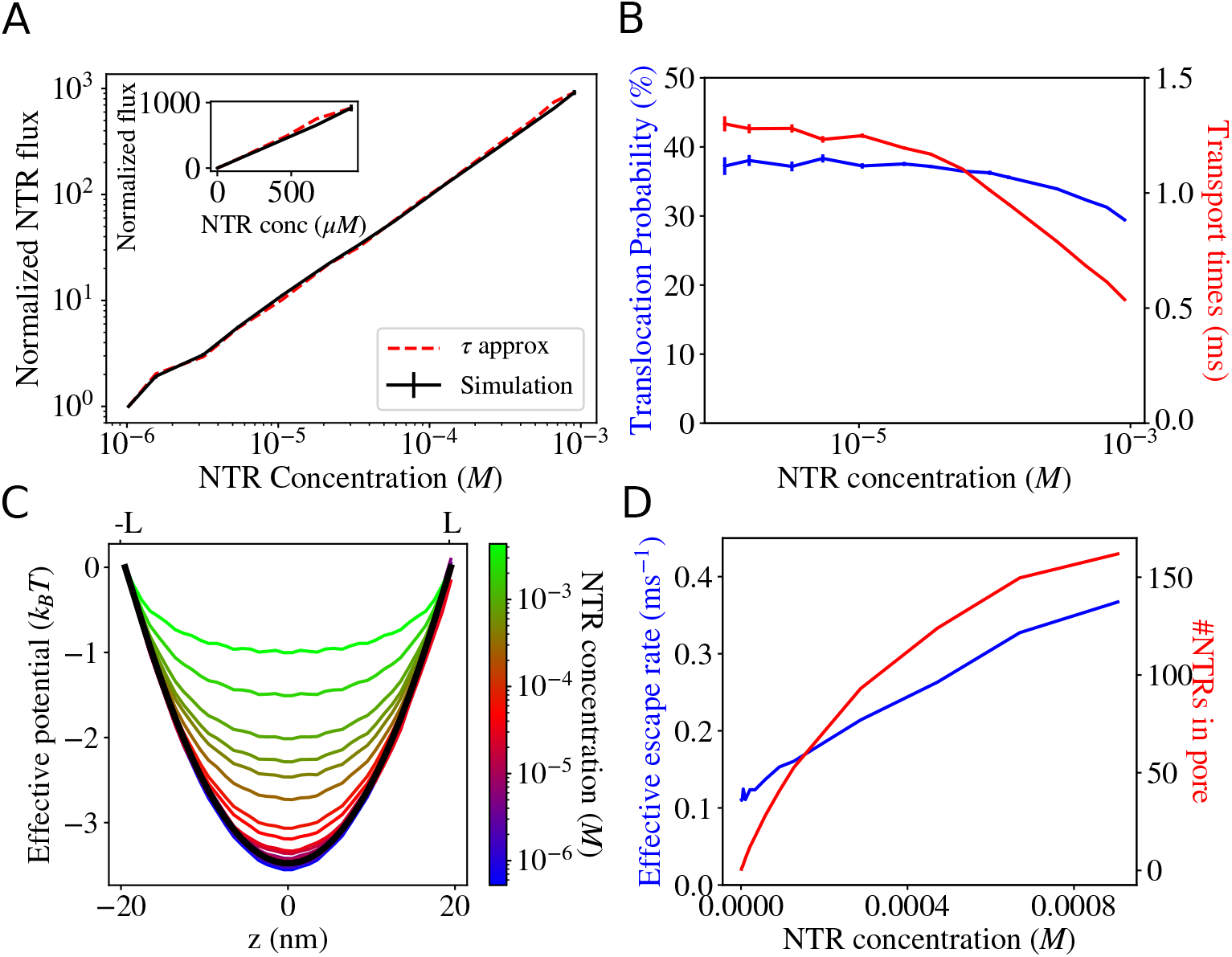
Competition-induced release observed in a pore where explicit FG nups are replaced by an implicit potential. (A) Flux through the pore (normalized by its value at an NTR concentration of 1 *µM*) does not saturate. Inset: flux plotted on linear axes. (B) Translocation probabilities and transport times both decrease as crowding within the pore increases. Error bars indicate SE. (C) Effective potentials measured from simulations. Crowding reduces the depth of the effective potential. Black line: the spatially-varying effective potential experienced by an NTR in the absence of crowding (see text for explanation). (D) While *N*_pore_, the average number of NTRs within the pore saturates (red line), see Equation 7, the flux does not due to a crowding-induced increase in the escape rates (blue line); see text for discussion.

As seen in Figure 4C, increased competition for space within the pore decreased the depth of the effective potential. Qualitatively, crowding therefore results in shorter transport times because the thermal fluctuations induced escape of the NTRs from the effective potential well is more likely to happen in shorter times in the more shallow potentials observed at higher concentrations [1, 58, 62]. The accompanying lack of flux saturation can be understood through a simple heuristic approximation for the flux (see Supplementary Information Section 1D):

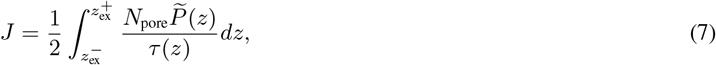

where 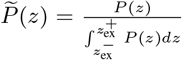 and 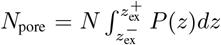 so that 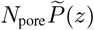 is again the number of NTRs within a “slice” *dz* of the pore, and 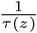 is the rate with which NTRs from this “slice” escape the pore (see Supplementary Information Section 1D for details). This heuristic approximation captures the behaviour of the normalized flux well as shown in Figure 4 and 2 (“*τ* approximation” lines). As shown in Figure 4D, while the number of NTRs within the pore, *N*_pore_, saturates at high NTR concentration, as expected, the average escape rate 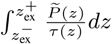 keeps increasing due to crowding induced shortening of the escape times. This *competition-induced release* is responsible for the observed lack of flux saturation even at high concentrations. This effect is expected to be rather general and independent of the precise molecular nature of the pore constituents.

However, as shown by Figure 4B, simple competition for space between NTRs in an effective potential is not sufficient to produce optimal single-molecule transport, where translocation probabilities increase with crowding (Figure 2D, shaded region). We show in the next subsections that the increase of translocation probabilities arises from the specific features of the heterogeneity of the FG nup distribution within the pore.

### 2.3 FG nup rearrangements induced by crowding are necessary for optimal single-molecule transport

To understand the origin of the unexpected regime of optimal single-molecule transport (Figure 2D, blue shaded region), we note that the crowding in the low NTR concentration regime not only decreases the average depth of the effective potential well (which, by itself, would result in decreased translocation probabilities) but also increases the steepness of the potential well at the pore entrance, as shown in Figure 3B (transition from blue to red lines).

The increase in the steepness of the potential near the pore entrance derives from changes in the compactness of the FG assembly at the pore peripheries. At low NTR concentrations, increasing the number of NTRs within the pore effectively helps “glue” the low cohesion FG nups into a more compact conformation which is pulled into the pore (Figure 5A to B) [11, 52]. This increases the density of FG nups in the region just inside the pore entrance (around *z* = −*L* + *R*), deepening the potential in this region via the increase in the density of the NTR binding sites (Figure 3B, blue to red). On the other hand, the density of the FG nups outside the pore decreases which leads to the increase in the effective potential outside the pore.

**Figure 5:**
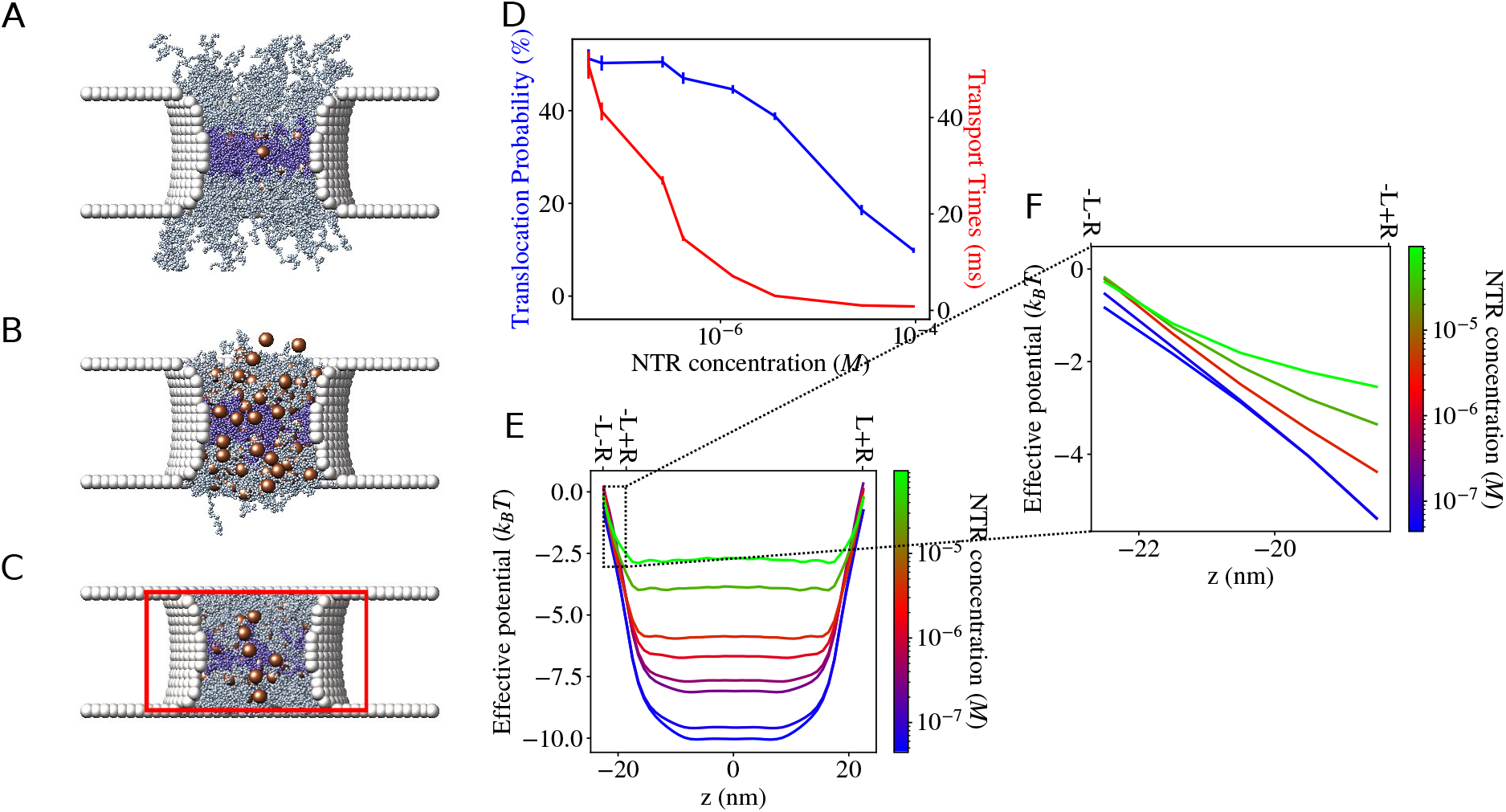
Effect of crowding-induced FG nup rearrangements around the pore exits. (A) Sample snapshot of our simulation at low NTR concentration: peripheral FG nups partially extend into the cytoplasmic compartment. (B) Sample snapshot of our simulation at a high NTR concentration: FG nups are mostly contained within the pore. (C) Sample snapshot of simulations with all FG nups confined within the pore by virtual “barriers” (red), thereby preventing FG nup rearrangements around the pore openings. (D) Translocation probabilities and transport times for the NPC model in panel C where FG nups are prevented from extending out of the pore. Without FG nup rearrangement, the translocation probability monotonically decreases with increasing crowding. Error bars indicate SE. (E) Effective potentials measured in the simulation where FG nups are artificially prevented from extending out of the pore. (F) Zoomed in effective potentials for representative NTR concentrations in the region from *z* = −*L* − *R* to *z* = −*L* + *R* (the bounds of the integral in the numerator of Equation 3). In contrast to Figure 3B, crowding monotonically decreases the slope of the potential in the region of the pore entrance.

Together, these effects result in the increasing translocation probabilities as follows from Equation 3. Intuitively, a steeper potential near the entrance reduces the probability of an NTR to diffuse backward and make an abortive exit.

At high NTR concentrations, after the compaction transition has already taken place, increasing the number of NTRs within the pore no longer changes the density of FG nups around the pore entrance. With FG nup re-arrangements no longer taking place, crowding increases the potential everywhere in the pore region, and the steepness of the potential close to the pore entrance again decreases (Figure 3B transition from red to green lines). Thus, the increase in the translocation probabilities at low NTR concentrations (Figure 2D, shaded blue box) is followed by the more expected decrease in transport probabilities at higher NTR concentrations.

To determine whether this phenomenon is responsible for the appearance of the regime where translocation probabilities increase with NTR concentration, we repeated the NPC simulations with additional virtual “barriers” that kept FG nups compacted within the pore for all NTR concentrations, without affecting the NTRs (Figure 5C). As shown in Figure 5E and F, without the changes to the compactness of the FG assembly, the steepness of the potential around the pore entrance does not increase with crowding, and correspondingly the translocation probability does not increase (Figure 5D). The ability of the low cohesiveness FG nups to rearrange around the NPC openings are therefore necessary for the regime of optimal single-molecule transport.

### 2.4 Spatial heterogeneity of the FG nup assembly is necessary for optimal single-molecule transport

Our minimal model of the NPC is based on extensive experimental evidence that the FG assembly contains at least two spatially and physically distinct regions: a central barrier region with a high density of relatively cohesive FG nups, and the peripheral “vestibule” regions with a relatively low density of less cohesive FG nups extending somewhat into the nucleus or cytoplasm [10, 14–18, 63, 64]. The previous subsection suggests that the crowding induced re-arrangements of the less cohesive vestibule nups around the pore exits are a key requirement for optimal single-molecule transport.

In this section, we therefore investigate whether the spatially heterogeneous architecture that includes both barrier and vestibule regions is necessary for producing this regime. To this end, we examined two variants of our NPC mimic that contain only one type of FG nup (either weakly or strongly cohesive). In the first variant, the cohesiveness of all FG nups is set to the same value as the (low) cohesiveness of the original vestibule FG nups. In the second variant the cohesiveness of all FG nups is set to that of the (high) cohesiveness of the original central barrier FG nups. The results of the high-cohesion case are shown in Figure 6. The more complex low-cohesion case is presented in Supplementary Information Section 1E. Notably, both variants with only one type of FG nup fail to produce the regime of the optimal single-molecule transport.

**Figure 6:**
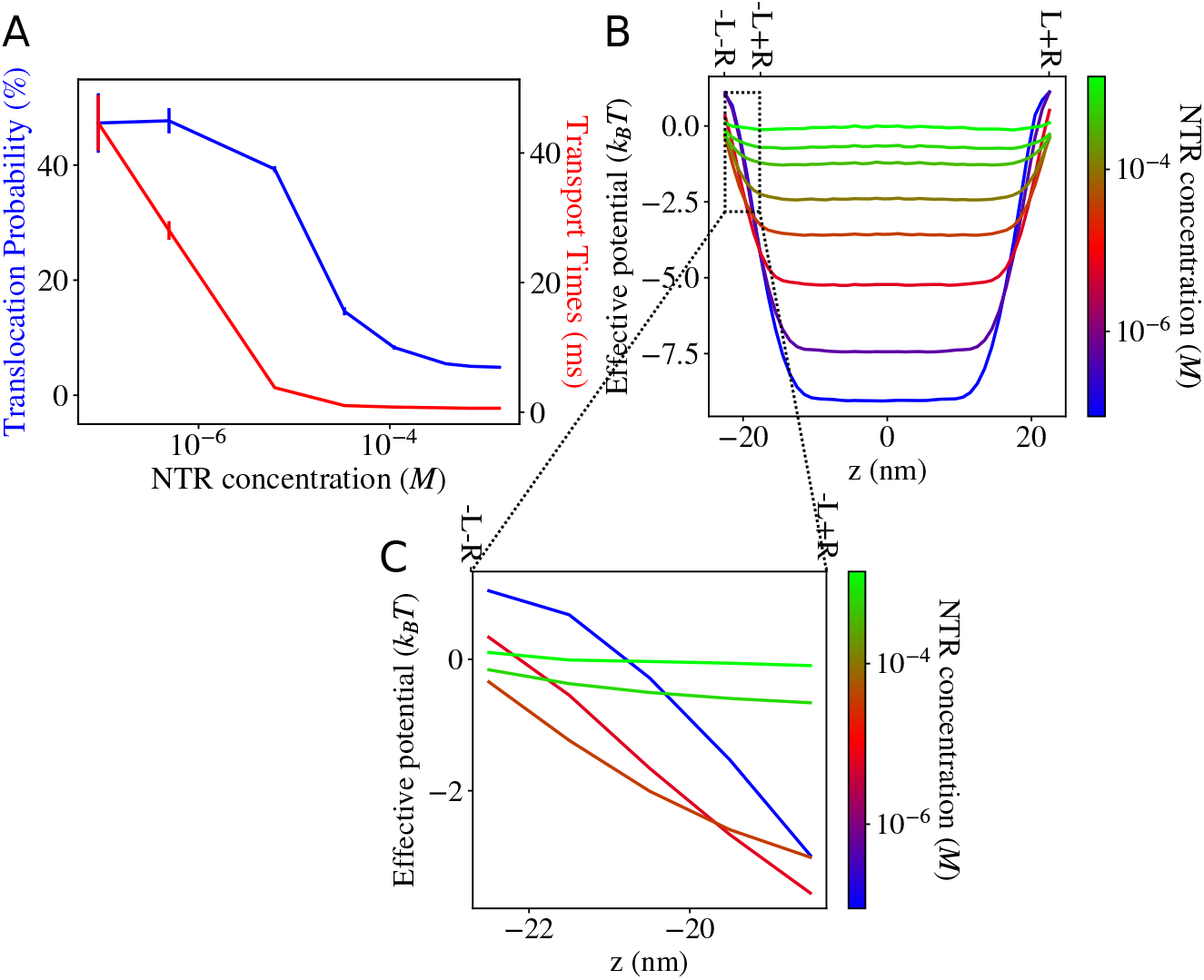
Transport through an NPC model with high cohesion nups only. (A) Both the translocation probability and the transport times for an NPC filled with FG nups with high cohesiveness decrease with crowding. Error bars show SE. (B) Effective potentials of NPC model with high cohesion nups only. (C) The steepness of the potential in the region around the pore entrance decreases monotonically with crowding.

As shown in (Figure 6), the NPC with only strongly cohesive FG nups does not reproduce optimal single-molecule transport, similar to the model where FG nups were replaced by an externally applied potential (Figure 4) and the model where the FG nups were forcibly contained within the pore (Figure 5). This occurs because the highly cohesive peripheral FG nups are already collapsed within the pore even in the absence of NTRs, and do not rearrange upon changes to the NTR concentration.

An architecture that combines the more cohesive FG nups in the centre of the pore with the less cohesive FG nups at the peripheries is therefore necessary for producing a regime of optimal single-molecule transport, where the translocation probabilities increase due to crowding while the transport times decrease.

### 2.5 Competition-induced release results in the appearance of slow and fast kinetics in the escape of NTRs from FG nup assemblies

The puzzle of transport times that decrease with crowding has been hypothesized to be connected to another long-standing puzzle in the field: the presence of a population of NTRs in FG nup assemblies with residence times much longer (minutes to hours) than typical transport times (milliseconds) [32, 36, 37, 40]. It has been suggested that the “slow” and the “fast” NTR phases have distinct functional roles whereby the strongly bound “slow” NTRs strengthen the FG nup barrier, while the “fast” NTRs rapidly transport cargoes through the NPC [32].

In this section we show that the existence of the apparent“slow” and “fast” NTR populations is a direct consequence of the competition-induced release observed in our model, discussed in Section 2B. However, we show that the “slow” and “fast” phases of NTRs are kinetically interchangeable populations rather than distinct spatially or functionally segregated ones.

As NTRs escape from an FG nup assembly, decreasing the crowding inside, the effective potential experienced by the remaining NTRs becomes deeper (similar to Figure 3A). Therefore, the NTR release times increase as fewer NTRs remain in the FG nup assembly. To test whether this effect can produce the appearance of “slow” and “fast” populations, we simulated our NPC model with absorbing boundaries on both ends of the simulation box. This allowed NTRs to irreversibly disappear into “bulk solution”, thereby mimicking the effect of rinsing permeabilized cells or surface-grafted assemblies with buffer.

We found that the decay of the number of NTRs remaining in the pore did not follow mono-exponential decay (Figure 7A), indicating the presence of more than one escape timescale. Instead, the decay is well approximated as a bi-exponential, consistent with the experimental reports of two populations of NTRs with “slow” and “fast” release timescales [32,36–40]. The apparent clear separation between the “slow” and the “fast” regimes occurs because the deepening of the effective potential due to competition-induced release (Figure 7B) proceeds at a non-uniform rate: rapidly initially (as many NTRs quickly escape in the initial stages of decay), and then very slowly (as it takes the remaining NTRs longer and longer to escape from the pore). This results in the appearance of a single “fast” timescale at the early time, while the deep potentials during the later part of the simulation determine the second “slow” timescale.

**Figure 7:**
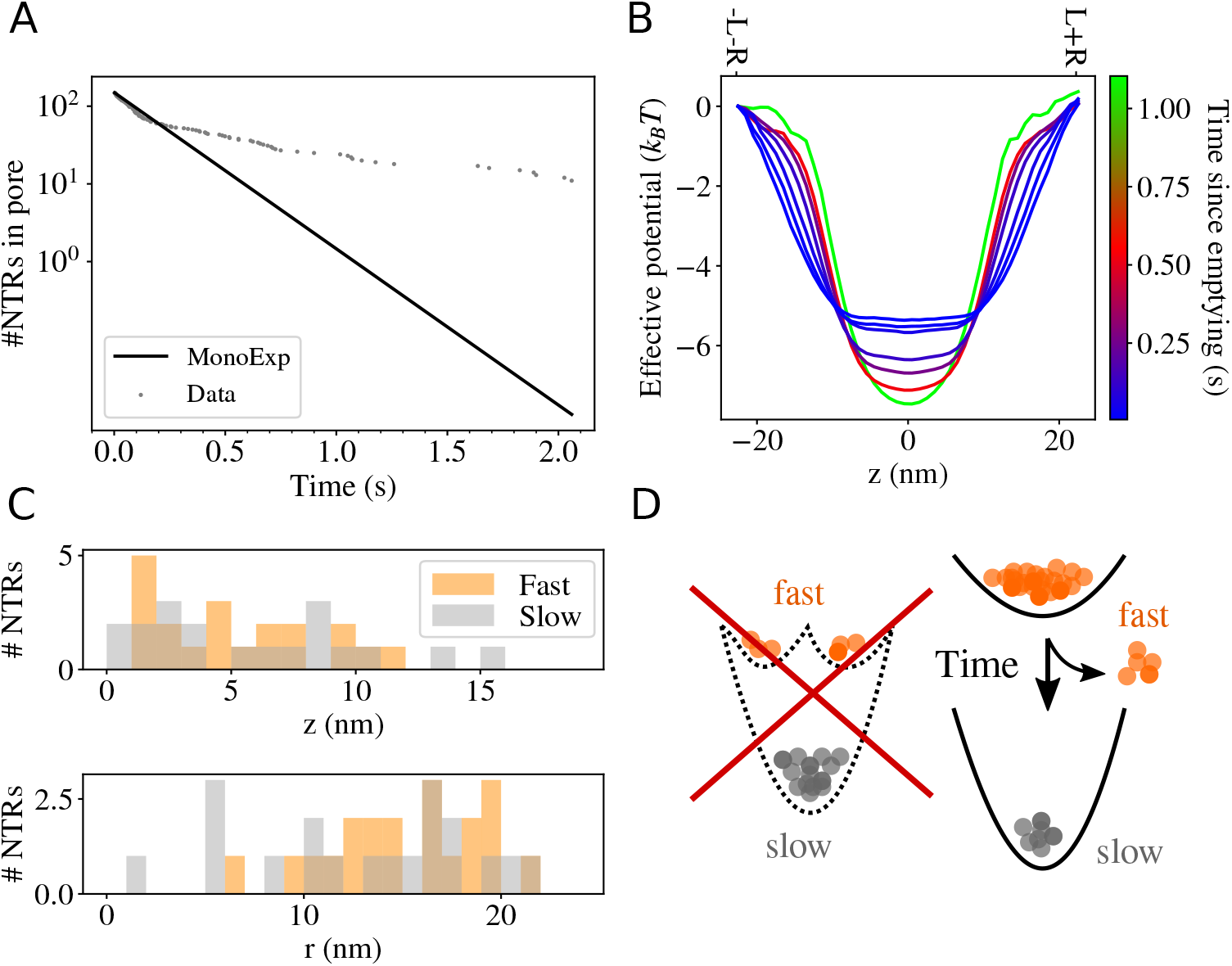
Analysis of the bi-phasic escape from the pore. (A) Grey dots indicate the number of NTRs remaining inside the pore at various times throughout our simulation. While the majority of the NTRs escaped within 0.1s, the pore was still populated by NTRs after 2s. The black line is a single-exponential fit to the early “fast” phase of the decay process. (B) The effective potential experienced by the NTRs inside the pore deepens as more NTRs escape due to competition-induced release, therefore increasing the dwell times as the pore empties. (C) Comparison of the initial *t* = 0 locations within the pore of the first 20 (“fast”) and the last 20 (“slow”) NTRs to leave the pore during the decay process (indicated in orange and grey colors respectively). (Top): initial position along the axial *z* coordinate. (Bottom): initial position along the radial *r* coordinate. (D) The average effective potential could be experienced by all NTRs, or it could be a composite of separate shallow and deep potentials (left), producing distinct fast and slow NTR populations. The results of our analysis are consistent with the scenario where the average potential is experienced by all NTRs (right), and fast and slow NTR populations are not distinct, but formed dynamically as the NTR population within the pore decays.

We next probed whether the NTRs in the long-bound fraction form a separate population, distinct from the NTRs that dissociate rapidly from the pore. Assigning a unique ID to every NTR in the simulation, we identified the first 20 NTRs to leave the pore (forming the “fast” group), and the last 20 NTRs to leave the pore (forming the “slow” group). Figure 7C shows the initial positions within the pore of the NTRs both for the “fast” and the “slow” groups, suggesting that the spatial regions occupied by these groups largely overlap, and the two groups are not spatially segregated.

To confirm this hypothesis, we ran several ensemble realizations of the system simulation starting from the same initial conditions but with different random trajectories (see Supplementary Information Section 1F for details). We found that the NTRs which we identified in the original realization as belonging to the “fast” group were not more likely to escape early in other ensemble trajectories.

In summary, we found no evidence of intrinsic – spatial or kinetic – differences between the “fast” and “slow” NTR populations.

## 3 Discussion

How the NPC is capable of simultaneously transporting large numbers of cargoes efficiently and rapidly has been a long-standing puzzle. While the ability to simultaneously transport multiple cargoes increases NPC throughput, the resulting crowding of the NPC channel was expected to present an obstacle to efficient and rapid transport [1, 8, 34, 65]. However, past experiments have suggested that crowding the NPC with NTRs or NTR-cargo complexes does not compromise transport. Rather, these experiments showed that the flux transmitted through the NPC does not visibly saturate even when the NTR concentration in the cytoplasm is an order of magnitude higher than the biological concentrations or the typical NTR-FG nup equilibrium dissociation constants [30, 31]. Even more surprisingly, single molecule experiments showed that increased crowding may lead to higher probabilities of making a successful translocation and shorter transport times [26] – a regime which we term *optimal single-molecule transport*. We provide a theoretical explanation of these results through simulations and theoretical analysis of a minimal NPC mimic with reduced structural complexity that includes only the main consensus features and components of the NPC transport system. We show that the model produces both the absence of flux saturation and a regime of optimal single-molecule transport.

Furthermore, we find that the non-saturation of flux is a rather general phenomenon that requires only competition between NTRs for space in an attractive potential well inside the NPC channel. Increased steric repulsion between the NTRs reduces the depth of the effective potential experienced by each NTR, thereby leading to faster transport and escape times – an effect we denote as *competition-induced release*. This effect also ensures the lack of saturation in the particle flux even though the number of NTRs within the pore saturates towards its maximal occupancy at high concentrations. In a system containing FG nups, we expect NTRs to compete for binding sites on FG nups in addition to competing for space, and coarse-grained theoretical calculations suggest that the effect of competition-induced release persists in this case [44].

Unlike the very general nature of competition-induced release, optimal single-molecule transport requires a particular spatial architecture of the FG nup assembly in the pore – both the central “barrier” region with more cohesive FG nups and the peripheral “vestibule” regions with less cohesive FG nups. The lower cohesion of vestibule nups allows them to re-arrange and compact around the pore openings in response to crowding, which is necessary for producing the increasing translocation probabilities in the optimal single-molecule transport regime. Accordingly, in the absence of the low cohesion vestibule nups, transport probabilities decrease monotonically with crowding. Similarly, without the higher cohesion central barrier FG nups, we also find no regime of optimal single-molecule transport where crowding improves both the speed and efficiency of transport.

The architecture incorporating both types of FG nups has long been suggested by observations [10, 13, 14, 39], and is likely to be evolutionarily conserved from yeast to mammalian cells [13,66]. It has been proposed that the low density/low cohesion “vestibules” at the pore periphery enhance the capture of NTRs while the central more cohesive FG nups serve as the “permeability barrier” [45, 67]. Our results suggest another potential functional purpose of this architecture as it allows the NPC to take advantage of crowding to achieve more efficient transport on the single-molecule level. Future work will explore the salience of these results on the background of the additional crowding in the nucleus and cytoplasm in live cells, and the presence of the nuclear basket and cytoplasmic filament structures that might shape FG assembly structure and dynamic at the pore exits, as well as other functional purposes selected for by evolution.

Somewhat surprisingly, we find that in our model the effects of crowding on the diffusion coefficients had no role to play in the clogging-free transport. In contrast to what may be expected from previous studies [68], we observe no increase of the diffusion coefficient within the pore due to crowding. These differences likely stem from the fact that the referenced study used a large colloidal particle coated with NTRs as a probe on an FG nup covered surface, and the observed decrease in the diffusion coefficient of the colloidal particle would have depended both on the NTR dissociation rate and diffusion coefficients on the surface of the FG polymer brush [68]. This was therefore a very different system to ours, where we measure the diffusion coefficients of NTRs within the FG assembly. See also Supplementary Information Section 1B.

Furthermore, the mechanism of competition-induced release allows us to explain previous observations of apparent “fast” and “slow” fractions of NTRs within the NPC [36, 37] and in vitro FG nup assemblies and artificial NPC mimics [32, 40]. As NTRs escape from an FG nup assembly, reduced competition between the remaining NTRs deepens the effective potential within the assembly, leading to very long escape times. Our analysis further shows that the long-lived fraction is likely not a separate population with *a priori* longer escape times, but is rather formed dynamically and stochastically as the earlier NTRs escape the pore.

One common thread emerging from our results is that competition-induced release is a powerful mechanism that can compensate for several potentially detrimental side effects of strong selectivity which requires relatively strong interactions of NTRs with the pore. Competition-induced release contributes to the explanation of the paradox of high selectivity versus the speed and throughput of the NPC. Contrary to the expectation that high thermodynamic selectivity would lead to the slowing down of transport, we observe that the strong interactions that attract large numbers of NTRs into the FG nup assembly produce crowding which results in rapid transport times through the competition-induced release mechanism. This results in clogging-free flux that increases linearly with NTR concentration without saturation even for concentrations much higher than the equilibrium dissociation constant between NTRs and the NPC (see Supplementary Figure S2). Our results suggest that the architecture of the NPC including only a few minimal components enables NPC to self-regulate the nucleocytoplasmic transport to achieve high efficiency in highly crowded conditions.

The general crowding-relieving mechanisms of the NPC identified in this work using minimal models can be applied to artificial nanopore design to harness the benefits of crowding based on the same principles. Understanding the native crowding-relieving mechanisms of the NPC has the potential to further efforts in understanding diseases that have been reported to be linked to the clogging of the NPC transport system [2]. Furthermore, the agreement between the minimal model and an array of independent experimental observations indicates that the model captures the essential properties of the NPC transport, forming the basis for future work. Future work will examine how NPCs regulate crowding when the cargoes are simultaneously transported bi-directionally through the NPC, and how very large cargoes such as viral capsids and ribosomal subunits navigate the NPC interior.

## Acknowledgements

We thank C. Gu, L.K. Davis, S.M. Musser, B.W. Hoogenboom and R.Y.H. Lim for insightful discussions. AZ acknowledges the support from the National Science and Engineering Research Council of Canada (NSERC) through the Discovery Grant Program RGPIN-2022-04909 and resources provided by Compute Canada through RRG-4094 RAC Allocation.

