## Supplementary Information for "Self-regulation of the nuclear pore complex enables clogging-free crowded transport"

### Supplmentary Information for
### Self-regulation of the nuclear pore complex enables clogging-free crowded transport

Tiantian Zheng[1] and Anton Zilman[1,2,*]
1 Department of Physics, University of Toronto, 60 St George st, Toronto, M1M 2P7 Ontario, Canada
2 Institute for Biomedical Engineering, University of Toronto, 164 College St, Toronto, ON M5S 3G9 Ontario, Canada
*

#### 1 Results

##### 1.1 Choice of the start and end points of the translocation attempts: effects on the optimal single-molecule transport regime

Our choice of the locations of where NTRs are considered to have left the NPC (thereby terminating abortive or successful entry events) are informed by the single molecule tracking experiments of [1]. As their experiments used human NPCs, which are longer than the yeast NPCs we modelled our mimic on, we opted for definitions which were similar in motivation, rather than preserving the exact distances. Their translocation attempts terminated once the NTRs reached positions where they were no longer be expected to interact with the NPC (i.e. they were further from the nuclear envelope than the distance between the tip of the cytoplasmic filaments to the centre of the NPC). In our model, as we had no cytoplasmic filaments, we chose the point at which NTRs were considered to have left the pore to be where the entire volume of the NTR was outside of the pore. Using our pore dimensions, this placed our pore exits at $z = \pm 22.5$ nm.

In order to understand whether our results were robust to this choice, we computed the translocation probabilities and transport times using Equations 3 and 4 for different choices of the assumed NPC entry and exit locations of the NTRs. As shown in Figure 4D), the decrease of transport times due to crowding is a very robust phenomenon and is insensitive to changes in these definitions. By contrast, the increase of the probabilities with crowding is more sensitive to the entry/exit locations definitions because it depends on NTR-induced re-arrangements of the FG nup cloud near the pore entrances. Figure 4C shows that the choice for the locations of the exits can be shifted by 10 nm outward while maintaining this regime. However, as 99% of the FG nups in the model are contained within the region between $z = -40$ nm and $z = 40$ nm, shifting the exit locations beyond these values leads to the disappearance of the optimal single molecule transport regime.

We also examined the choice of the definition of the pore entry locations as the point where the entire volume of the NTR was inside the pore (at $z = -17.5$ nm). Figure 4A and B show that shifting the definition of the pore entrance towards the cytoplasmic exit does not affect the existence of the optimal single molecule transport regime.

##### 1.2 Measuring effective potentials and diffusion coefficients

We ran auxiliary simulations in order to measure the effective potentials and diffusion coefficients inside the pore at different NTR concentrations. To measure the equilibrium effective potential in the pore, we ran simulations where NTRs were allowed to equilibriate throughout the simulation box to estimate the equilibrium one-dimensional probability distribution $P_{\text{eq}(z)}$ (in contrast, the simulations in the main text were run with a non-zero steady-state flux). The effective potential in the pore was obtained as $U = -ln(P_{\text{eq}(z)})$ [2–4].

As the FG nup density may have an effect on the effective diffusion coefficient, we measured the diffusion coefficient separately in the low density peripheral and high density central regions of the pore (demarcated in Figure 5A). For each NTR concentration, the two regions of the pore and the equilibrium number of NTRs and FG nups contained within were placed in a separate simulation box with periodic boundary conditions, so that NTRs were effectively diffusing within an infinite, rugged pipe formed by repeats of this region (Figure 5B). Despite the complex environment within the pore, we observe normal diffusion of the NTRs in the pore as illustrated in Figure 6 that shows sample MSD plots obtained separately for the central and peripheral regions within the pore (see Methods), at various concentrations. The MSDs of the NTRs were then measured, and the effect of the spatially varying effective potential within each region of the pore was removed using the Zwanzig correction [5] to the diffusion coefficient:

$$D_{\text{true}} = \langle e^{U(z)} \rangle_z D_{\text{app}} \langle e^{-U(z)} \rangle_z$$

where $D_{\text{true}}$ is the true diffusion coefficient, $D_{\text{app}}$ is the apparent diffusion coefficient obtained from the MSD measurements, and $U(z)$ is the effective potential within the region of the pore.

#### 1.3 Changes in the diffusion coefficient due to crowding cannot explain the clogging-free response

Within the NPC model of this paper, due to the differences in cohesiveness between peripheral and central FG nups and non-uniform shape of the pore, the density of FG nups is non-uniform throughout the channel. This difference in FG nup density along the pore results in different effective diffusion coefficients at the center and at the peripheries of the pore. Within our single-particle 1D diffusion model, the effective diffusion coefficient $D(z)$ is therefore approximated as piecewise-constant:

$$D(z) = \begin{cases} D_{\text{out}}, & z < -L \\ D_{\text{in}}^{\text{P}}, & -L \leq z < -L^C \\ D_{\text{in}}^{\text{C}}, & -L^C \leq z < L^C \\ D_{\text{in}}^{\text{P}}, & L^C \leq z < L \\ D_{\text{out}}, & L \leq z \end{cases} \tag{1}$$

where $D_{\text{out}}$ is the diffusion coefficient outside the pore, $D_{\text{in}}^{\text{P}}$ is the diffusion coefficient in the peripheral regions of the pore, and $D_{\text{in}}^{\text{C}}$ is the diffusion coefficient in the central region of the pore. $L^C = 6.2$ nm is the boundary between the central and peripheral regions of the pore (see Supplementary Figure 5). $D_{\text{out}}$ is given by the Stokes-Einstein equation and has no dependence on crowding. Figure **??**C shows that both $D_{\text{in}}^{\text{P}}$ and $D_{\text{in}}^{\text{C}}$ are non-increasing as crowding increases.

We find that the non-increasing diffusion coefficients which we measure within the pore cannot produce by themselves any of the clogging free behaviours. Equation 6 shows that decreasing $D(z)$ would lead to a decrease in flux; therefore without the changes to the effective potential, we would expect to see a saturation of the flux due to crowding.

We further investigated whether the changes to the diffusion coefficient due to crowding alone could produce optimal single-molecule transport in the hypothetical case of an effective potential that did not change with crowding. Figure 7 shows that if the effective potential is fixed, the changes in the diffusion coefficient due to crowding do not produce optimal single-molecule transport. We conclude that the clogging-free regime we observe occurs due to the changes of the effective potential due to crowding rather than crowding effects on the diffusion coefficients.

#### 1.4 A heuristic model for the absence of the flux saturation with crowding

Assuming that the NTR distribution within the pore is almost symmetric (which is true even at nonequilibrium steady states, see Supplementary Figure 3B), and therefore approximately half of the total flux leaving the NPC exits into the nucleus. The flux can be heuristically approximated as

$$J = \frac{1}{2} \int_{z_{\text{ex}}^-}^{z_{\text{ex}}^+} \frac{N_{\text{pore}} \widetilde{P}(z)}{\tau(z)} dz \tag{2}$$

where $\frac{1}{\tau(z)}$ is the average rate with which NTRs starting from an initial position $z$ escape from the NPC (in either direction). $N_{\text{pore}}$ is the total number of NTRs within the pore, and $\widetilde{P}(z) = \frac{P(z)}{\int_{z_{\text{ex}}^-}^{z_{\text{ex}}^+} P(z)}$ is the 1D probability density of NTR positions within the pore, such that $N_{\text{pore}} \widetilde{P}(z)$ is the expected number of NTRs within a slice $dz$ of the pore. $\tau(z)$ is the average residence time, given as the mean first passage time to escape through either pore opening [6–8]:

$$\tau(z) = \int_{z}^{z_{\text{ex}}^+} \left[ \frac{e^{U(z')}}{D(z)} \int_0^{z'} e^{-U(z'')} dz'' \right] dz' \tag{3}$$

The results of this model are shown in the main text in Figures 2C and 4A ("$\tau$ approx" lines).

#### 1.5 An NPC model containing only low cohesion FG nups cannot produce optimal single-molecule transport

The case of an NPC filled with only low cohesion FG nups is somewhat more complicated than the NPC filled with only high cohesion FG nups. As shown in Figure 8A we observe two distinct regimes of the dependence of the translocation dynamics on crowding at low and high NTR concentrations. At low NTR concentrations both translocation probabilities and transport times increase with NTR concentration. The qualitative explanation for this effect is that due to the low cohesion of the FG nups the density of FG nups inside the pore is low in the absence of NTRs. As the number of NTRs inside the pore increases, their interaction with FG nups initially draws more nups into the pore, deepening the effective potential in the pore and leading to the increase in both translocation probabilities and transport. At high concentrations, the FG nups are already collapsed into the pore and the effect of competition between NTRs begins to dominate, making the effective potentials shallower as crowding increases. This leads to a decreasing translocation probability with crowding, and transport times that initially drop and then gradually increase due to jamming [8]. Notably, we do not observe and optimal single-molecule transport with increasing translocation probability and simultaneously decreasing transport times neither at the high nor at the low concentration regimes.

#### 1.6 Interchangeability of "slow" and "fast" NTRs within FG nup assemblies

In Section 2E we observed a biphasic decay of NTRs within the pore which was consistent with previous observations of "fast" and "slow" NTR populations [9–12].

We wanted to further understand whether the "fast" and "slow" groups were dynamically distinct populations predisposed to escaping at different rates, or if the composition of these groups was random.

To this end, we first identified the NTRs which were among the first 20 to leave the pore as the "fast" group, and the NTRs which had been among the last 20 to leave the pore as the "slow" group (out of the initial 150 NTRs in the pore). We then ran short simulations to sample 100 random realizations of the early stages of our emptying pore, using the same initial conditions and changing only the seed of the random number generator, and recorded the identities of the first five NTRs to escape each time.

We found that NTRs identified as members of the group of 20 "fast" NTRs and the group of 20 "slow" NTRs in the initial simulation were equally likely to be among the first five to leave in the alternative realizations of the simulation. This is quantified in Figure 9 which shows that the proportion of the alternate realizations where $n$ NTRs from the "fast" or "slow" groups were among the first 5 to escape agrees with the binomial distribution $\mathcal{B}(n, p)$ where $p$ is the probability of randomly selecting an NTR from the "fast" or "slow" groups by picking any NTR initially within the pore. Therefore NTRs from the originally identified "fast" group are not more likely to escape the pore first compared to NTRs which were not from this group. This therefore indicates that the "fast" and "slow" groups cannot be identified *a priori*, but are formed dynamically as the pore empties.

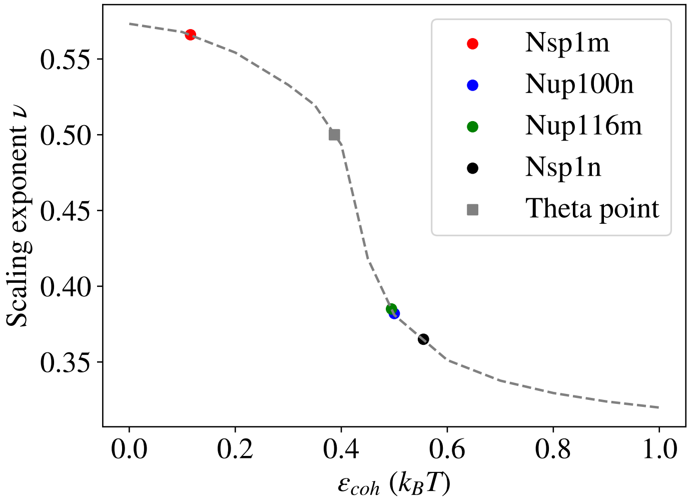


Supplementary Figure 1: FG nup cohesion calibration: The scaling parameter $\nu$ determines the average linear dimension of the polymer through $R \propto N^{\nu}$, quantified here as the radius of gyration. Here we show the experimentally determined scaling exponents of select FG nups [13, 14] and the corresponding $\epsilon_{\text{coh}}$ values in our simulation setup. The cohesiveness of the peripheral FG nups in our NPC model was $\epsilon_{\text{coh}} = 0.3\ k_B T$, and $\epsilon_{\text{coh}} = 0.5\ k_B T$ for the central FG nups. These values were chosen to lie on opposite sides of the coil-globule transition ("theta point"), reflecting the two types of conformations found in FG nups [13].

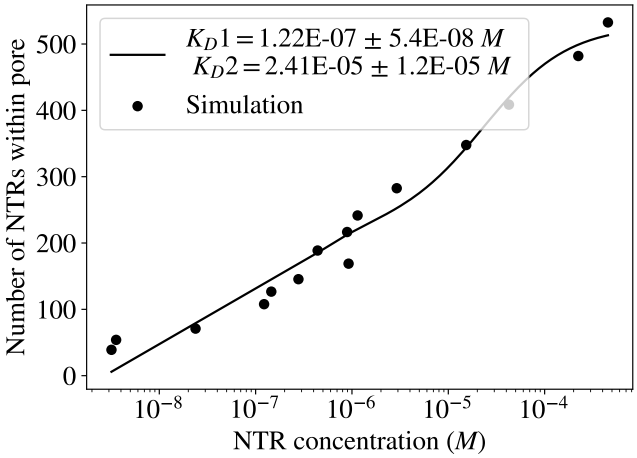


Supplementary Figure 2: Our NPC model displays strong selectivity for NTRs. The accumulation of NTRs in our NPC model is fit by a 2-component Langmuir model producing $K_{D_1} = 1.2 \times 10^{-7}$ $M$ and $K_{D_2} = 2.4 \times 10^{-5}$ $M$. These values agree well with experimentally measured values [15]. However, the bulk flux through our NPC model does not saturate even at concentrations well above both of these $K_D$ values.

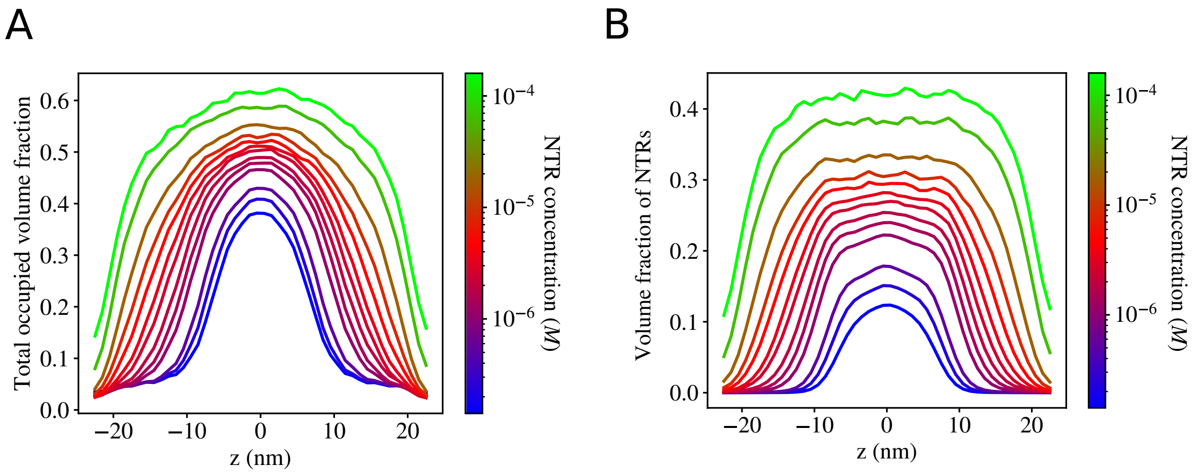


Supplementary Figure 3: Volume fractions of NTRs and FG nups within the pore. (A) The total volume fraction occupied by FG nups and NTRs within our simulation reaches more than 0.6 in parts of the pore, which is close to the close random packing densities of 0.62-0.64. Despite such a crowded interior, we observe no saturation of the flux through the NPC or increase in the transport times. (B) The distribution of NTRs within the pore at steady-state is almost symmetric for NTR concentrations, allowing us to make simplifying assumptions in the derivation of Equation 7.

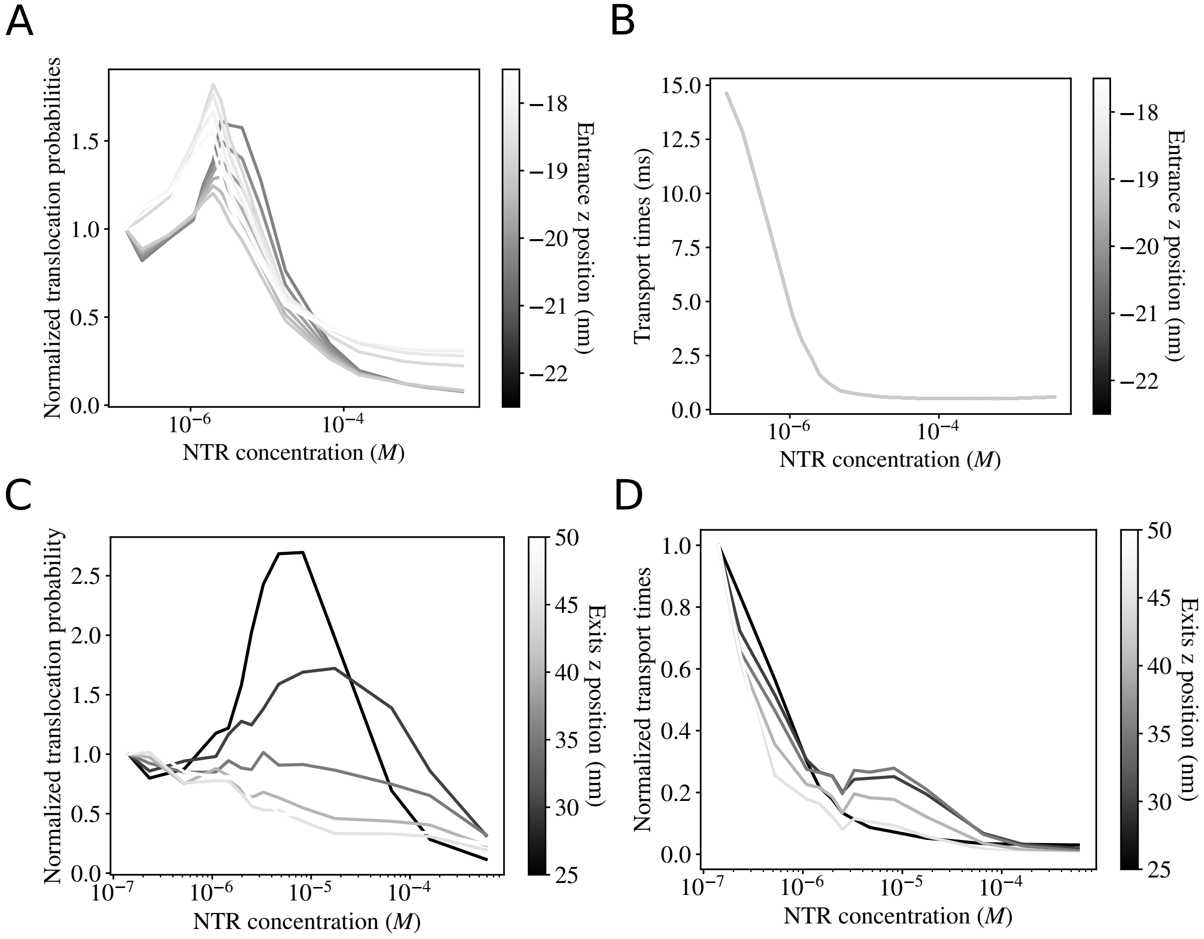


Supplementary Figure 4: Testing the alternative definitions of the beginning and end of a translocation attempt. The nuclear envelope extends between $z = \pm 20$ nm. (A) Translocation probabilities for different entrance positions, with values normalized by the translocation probability at the lowest NTR concentration. (B) Differences in the transport times using different choices of the entrance positions are negligible. (C) Translocation probabilities for different choices exit positions, with values normalized by the translocation probability at the lowest NTR concentration. The choice of exit positions is always symmetric such that $-z_{\mathrm{ex}}^{-} = z_{\mathrm{ex}}^{+}$. In all simulations an NTR was consideded to have entered the NPC when its center reached one diameter to the right of the cytoplasmic exit location. (D) Transport times for different choices of the exit positions, normalized by the times at the lowest NTR concentration.

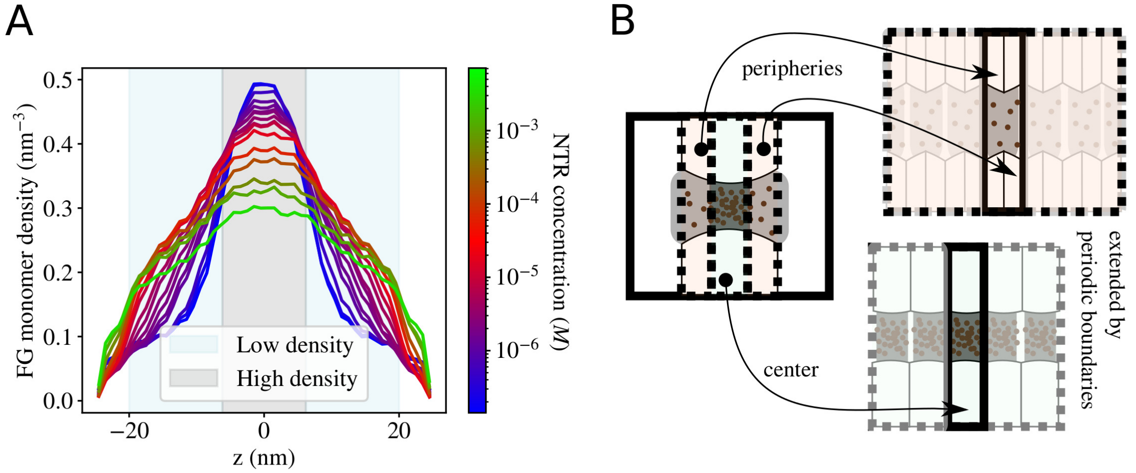


Supplementary Figure 5: The setup for measuring the effective diffusion coefficients from simulations. (A) To account for the differences in FG nup densities at different regions within the pore, the diffusion coefficients were measured separately for the central (grey) and peripheral (blue) regions of the pore. (B) For each NTR concentration, the central and peripheral regions of the pore (and the NTRs contained within) were each placed in a separate simulation box with periodic boundary conditions, and the MSDs of the NTRs were measured within these infinite, irregular channels.

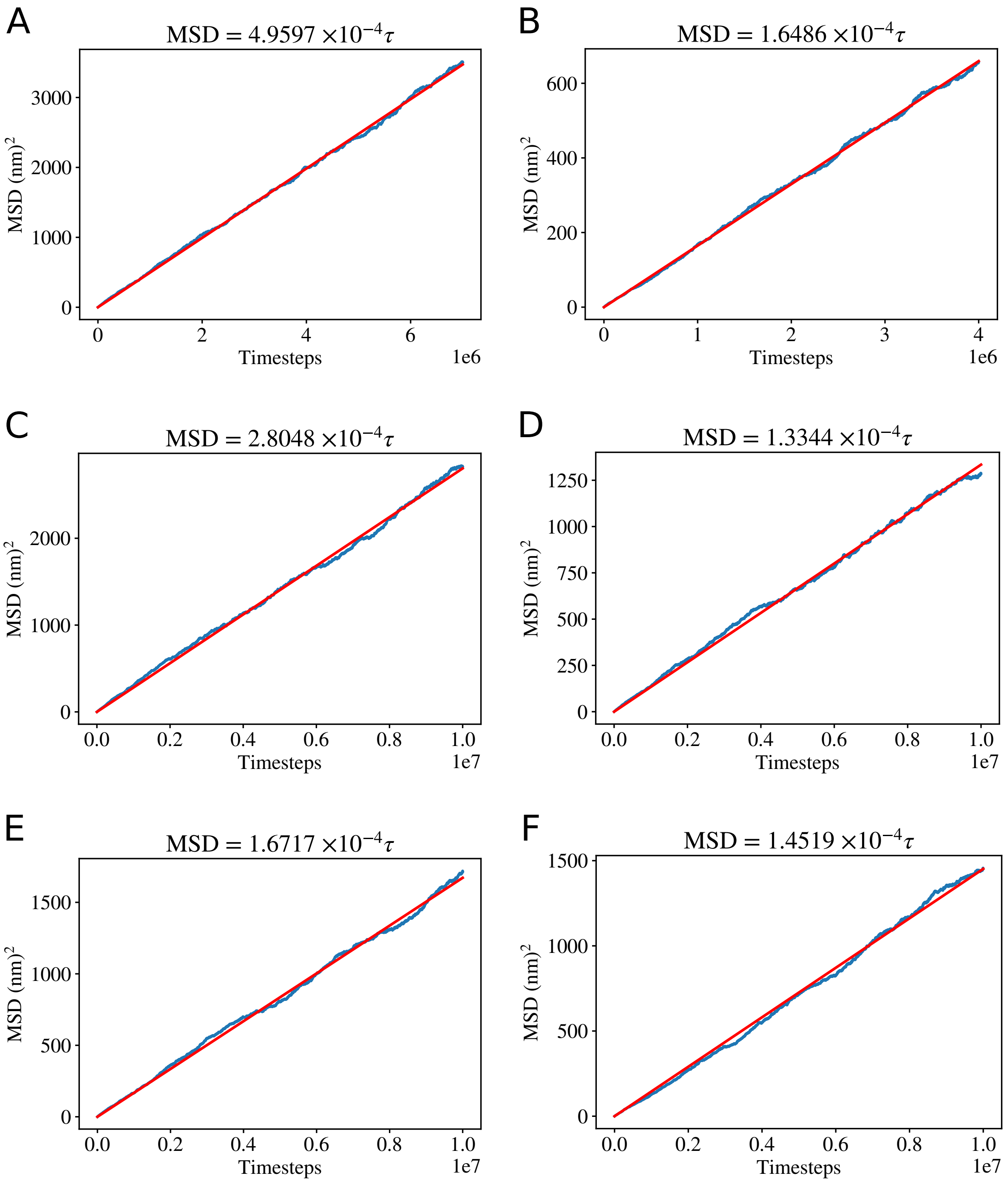


Supplementary Figure 6: Sample MSD measurements within the central and peripheral regions of the pore at different NTR concentrations. (A) MSD measured in central region of the pore, NTR concentration $= 1.42 \times 10^{-7} M$. (B) MSD measured in peripheral region of the pore, NTR concentration $= 1.42 \times 10^{-7} M$. (C) MSD measured in central region of the pore, NTR concentration $= 2.49 \times 10^{-6} M$. (D) MSD measured in peripheral region of the pore, NTR concentration $= 2.49 \times 10^{-6} M$. (E) MSD measured in central region of the pore, NTR concentration $= 1.60 \times 10^{-4} M$. (F) MSD measured in peripheral region of the pore, NTR concentration $= 1.60 \times 10^{-4} M$.

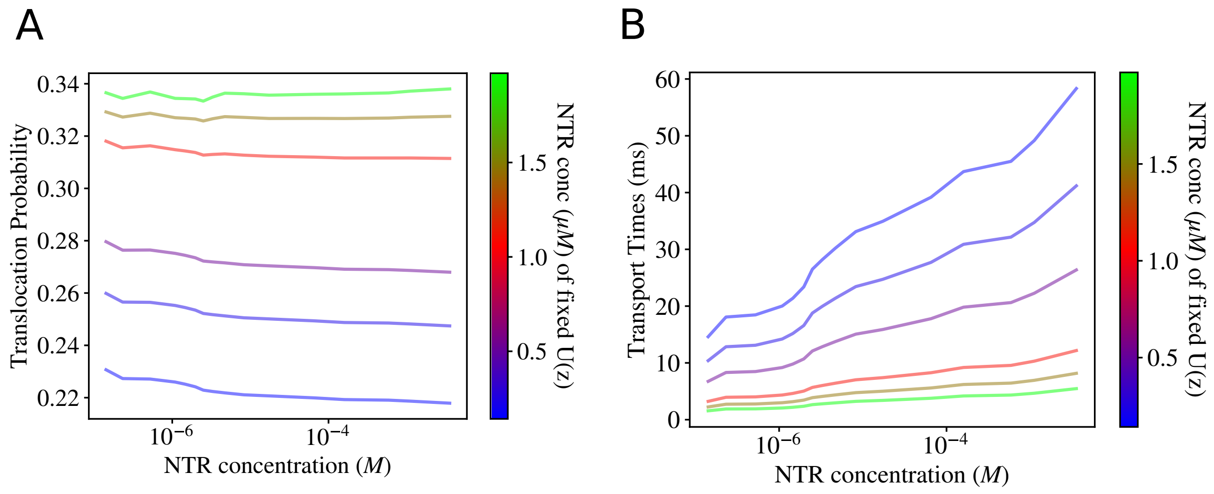


Supplementary Figure 7: The changes in the effective diffusion coefficient alone cannot produce increasing translocation probabilities or decreasing transport times. (A) We fix $U(z)$ to be the effective potential measured at various low NTR concentrations, and plot Equation 3 showing how the changes to the diffusion coefficients can only result in translocation probabilities which decrease with crowding when the effective potential is held fixed. (B) Similarly fixing $U(z)$ to be the effective potential measured at various NTR concentrations, and plotting Equation 4 shows that the changes to the diffusion coefficients can only result in transport times which increase with crowding.

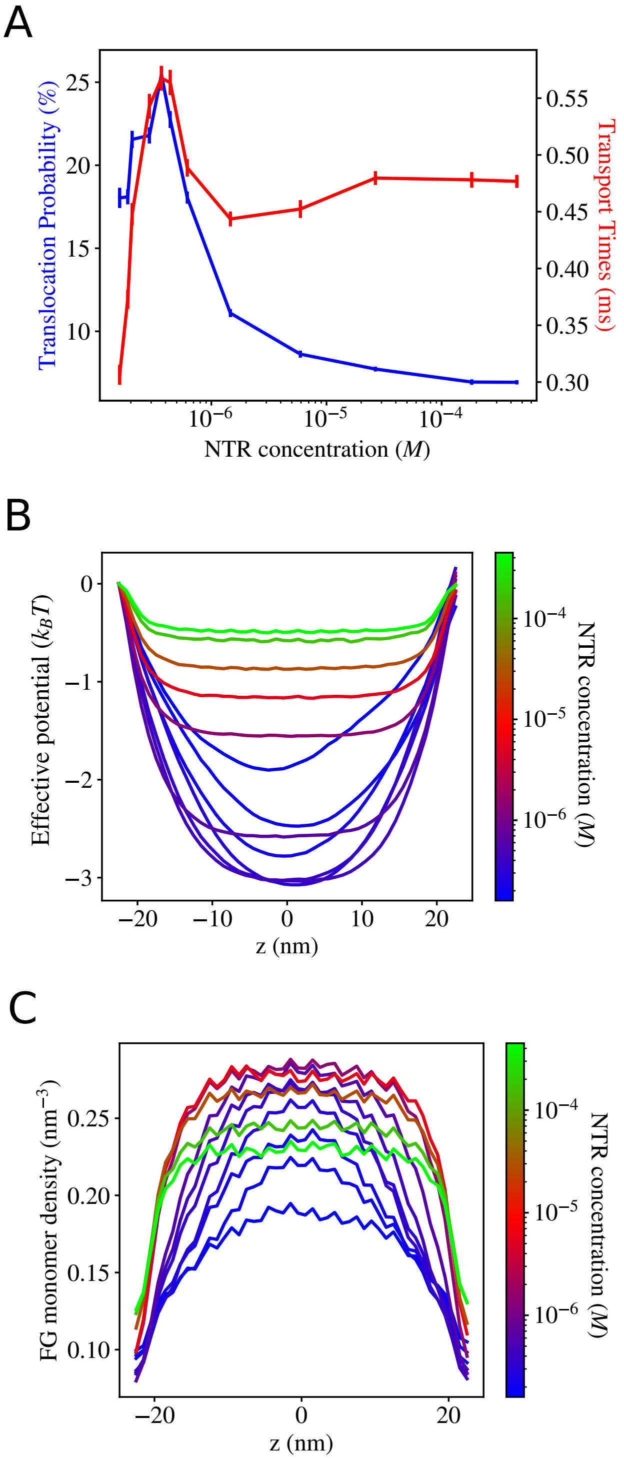


Supplementary Figure 8: Simulation results for an NPC filled with only low cohesion FG nups. (A) Translocation probabilities and transport times. (B) The effective potentials within the pore, relative to the potential at the cytoplasmic exit. As the NTR concentration increases, the effective potential deepens (blue to light purple), then becomes shallower (light purple to green). (C) As the NTR concentration increases, the density of the FG nups within the pore first increases (blue to purple), then decreases (purple to green) as they are squeezed out of the pore.

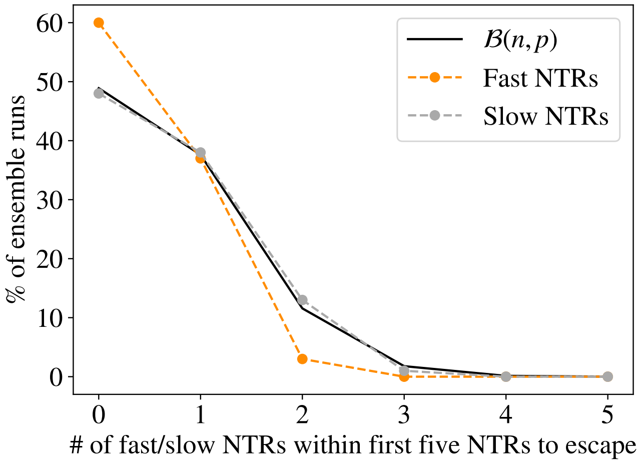


Supplementary Figure 9: Repeated sampling of the first five NTRs to leave the NPC yields similar distributions for NTRs from the original "fast" and "slow" groups identified in our initial, full-length simulation, see text. The proportion of the 100 ensemble runs where $n$ "fast" or "slow" NTRs were observed among the first five to escape is well approximated by the binomial distribution resulting from the hypothesis that all NTRs initially within the pore are equally likely to escape.
